## Supporting information for "Artificial soils reveal individual factor controls on microbial processes"

### Artificial soils reveal how individual matrix parameters control microbial processes

#### List of contents

1. Figures S1 - S12
2. Tables S1 - S3
3. Artificial Soil Production Protocol

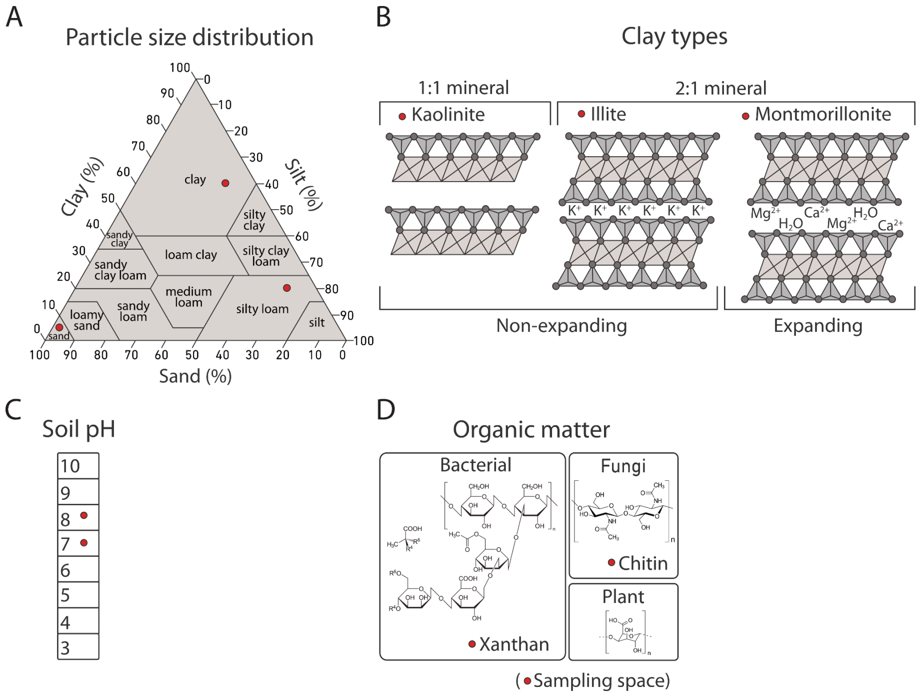


**Supplemental Figure 1. Sampling space for the artificial soils. (A)** The three particle size distributions (sand, silt loam, and clay) mixed to make artificial soils (red circles) are mapped onto a USDA soil texture triangle. **(B)** The different clay types are added at different levels to vary mineralogy. **(C)** The soil pH values are tuned by adding $CaCO_3$ or $Al_2(SO_4)_3$. **(D)** OM is added from different sources.

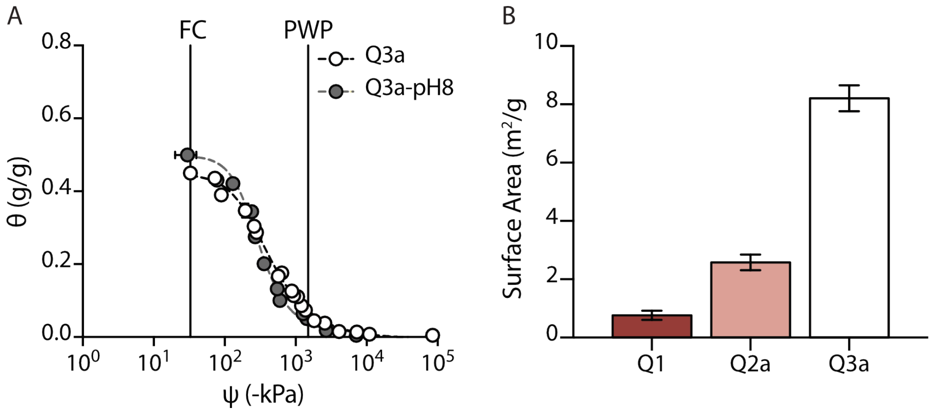


**Supplemental Figure 2. Additional artificial soil characteristics.** (A) Water retention curve of quartz-based soil with clay particle size distribution lacking (Q3a, white circles) and containing $CaCO_3$ (Q3a-pH8, gray circles). FC = field capacity, PWP = permanent wilting point. (B) Surface area of quartz-based artificial soils with different particle size distribution. Error bars represent one standard deviation calculated from 3 experiments.

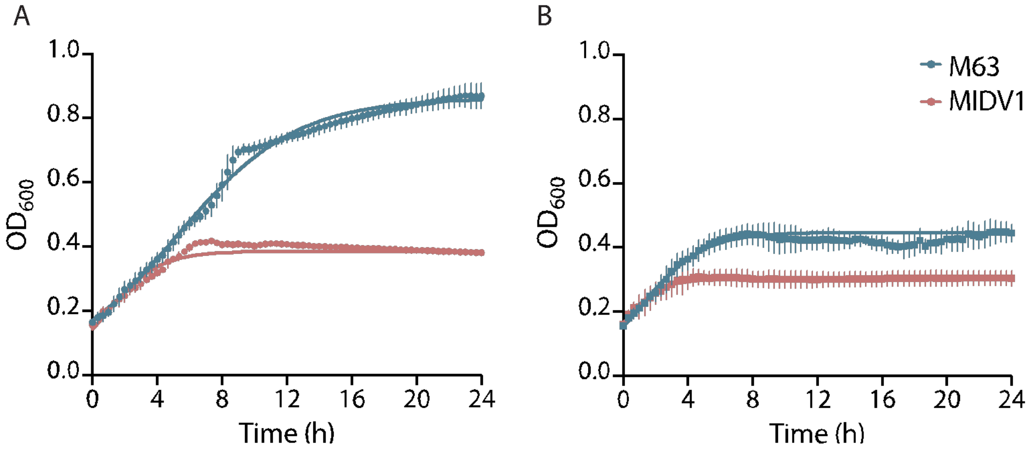


**Supplemental Figure 3. Microbial growth in environmentally-relevant media.** (A) Ec-MHT and (B) Bs-MHT growth (OD600) in M63 medium (blue) having an osmotic potential (1443 kPa) that exceeds natural soils and MIDV1 medium (red) having a potential (319 kPa) that is environmentally relevant. Experiments were performed in triplicate, with error bars representing one standard deviation. A fit of the Ec-MHT data to a logistic growth model (line) yielded different growth rates and delays in M63 medium (0.27 hours-1 and 3.69 hours) and MIDV1 (0.62 hours-1 and 1.62 hours). Different values were also obtained with Bs-MHT grown in M63 (0.53 hours-1 and 1.87 hours) and MIDV1 (0.87 hours-1 and 1.16 hours) medium.

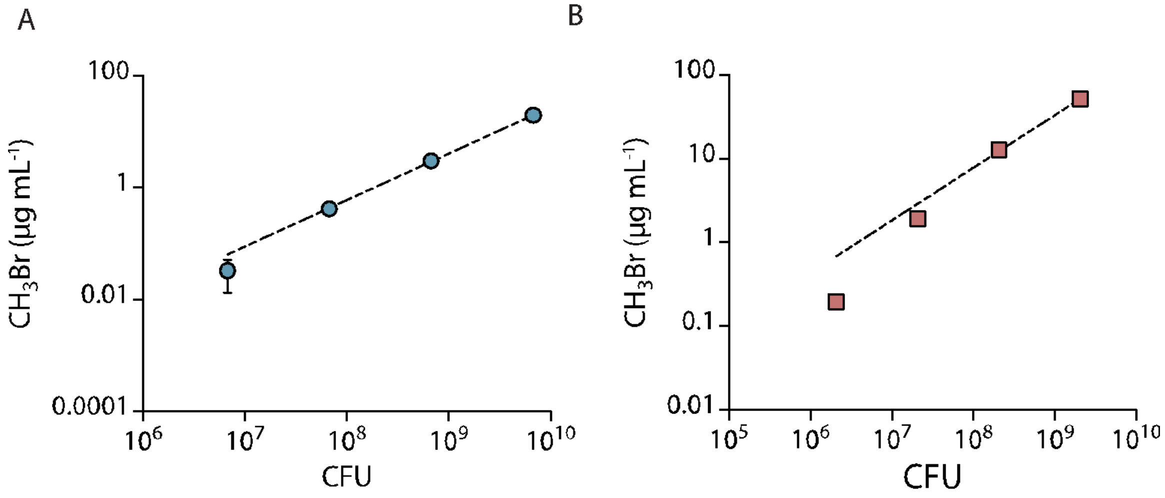


**Supplemental Figure 4. Relationship between $CH_3Br$ gas production and CFU.** Serial dilutions (10x) of (A) Ec-MHT and (B) Bs-MHT were added to 2 mL sealed glass vials containing 200 µL MIDV1 medium. Gas was measured using GC-MS following a 3-hour incubation at 30°C. To quantify CFU, cells were spread on LB-agar plates, incubated at 30°C for 24 hours, and visually inspected to count colonies. Gas production per CFU fitted to a log-log linear regression model. Both fits have R2 = 0.99. All experiments were performed in replicate, with error bars representing one standard deviation.

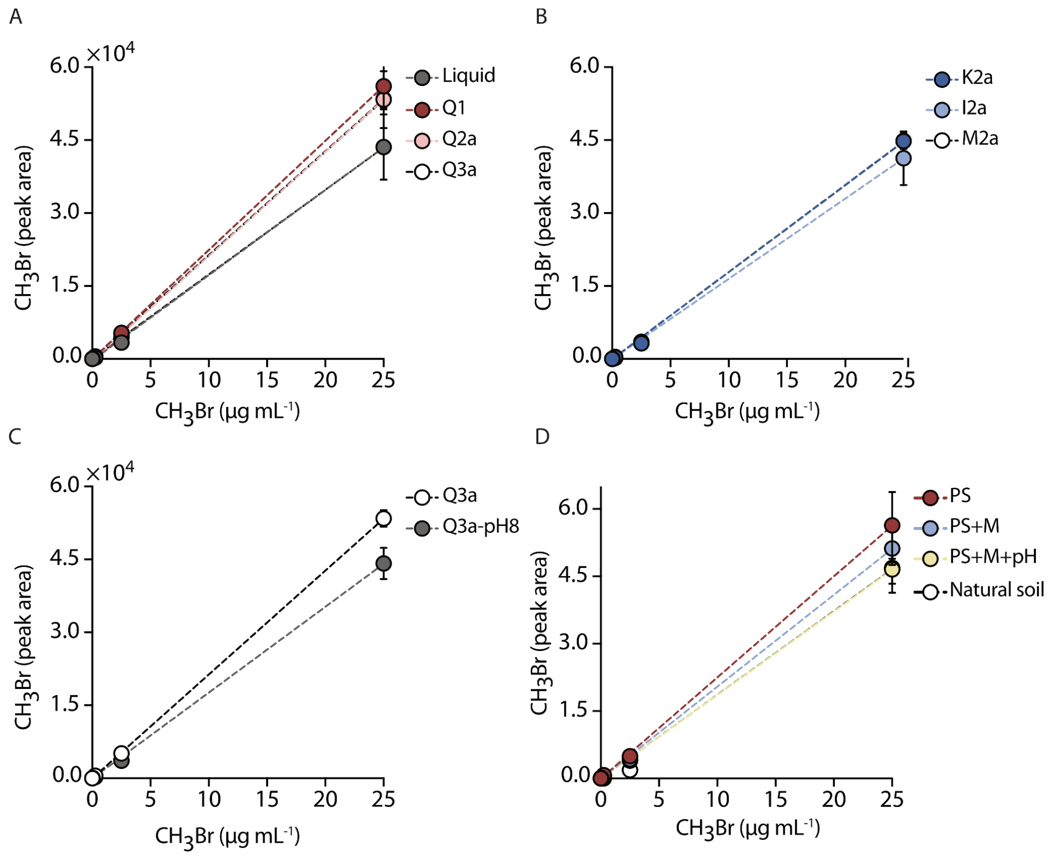


**Supplemental Figure 5. $CH_3Br$ standard curves in the artificial soils.** To obtain standard curves for calculating in experiments, we evaluated how artificial soils varied in their headspace gas following addition of defined amounts of indicator gas standards. Artificial soils were hydrated to the same water content across the soil types used to study each physicochemical property using MIDV1 medium. Serial dilutions (10x) of $CH_3Br$ chemical standard were added to artificial soils that vary in (A) particle size distribution, (B) mineralogy, and (C) pH. (D) Measurements were also performed in the Mollisol and the artificial soil series resembling the Mollisol. After capping, vials were incubated for 6 hours at 30°C, and headspace gas was measured using a GC-MS. The dashed lines indicate linear fits to the data. Error bar indicates one standard deviation calculated using an n=3.

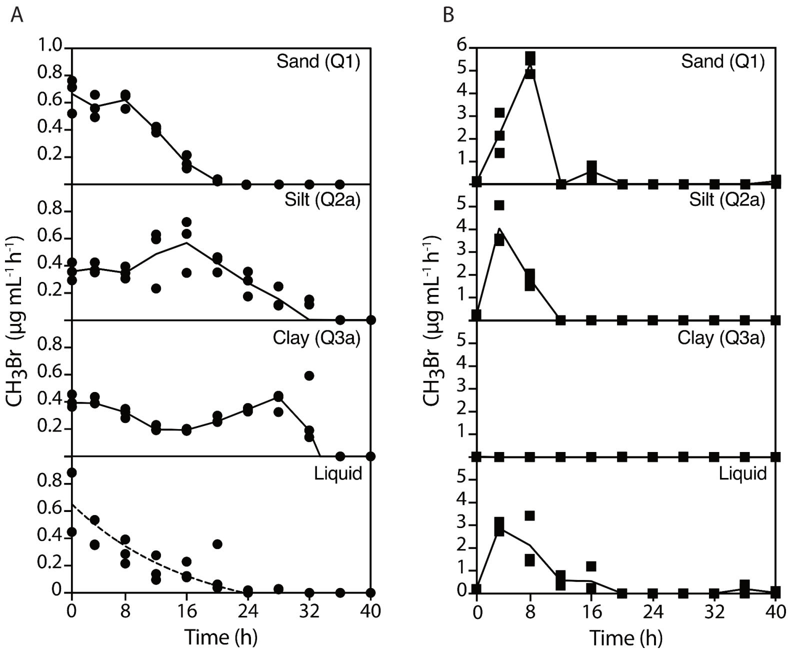


**Supplemental Figure 6. $CH_3Br$ production rates in soils having different particle sizes.** Effect of participle size distributions on dynamic $CH_3Br$ production with soils containing (A) Ec-MHT and (B) Bs-MHT. Each symbol indicates a single replicate. The solid lines represent interpolation using the mean calculated from three separate experiments. The dashed lines in liquid medium represent a fit to a single exponential decay model (R2= 0.71).

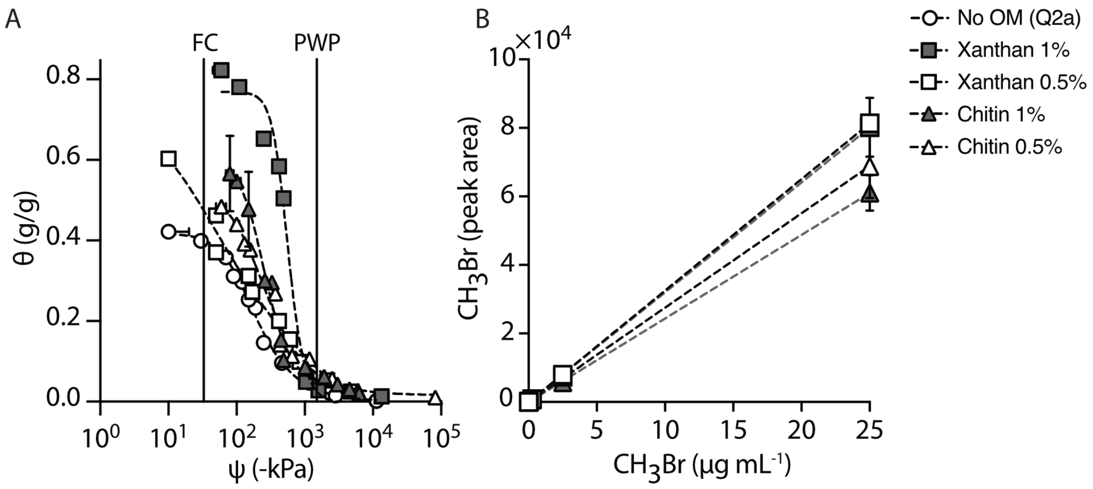


**Supplemental Figure 7. Characterization of artificial soils with OM.** (A) Soils that vary OM source and content present different water retention curves, including no OM (circle), xanthan gum (square) or chitin (triangle) at 0.5% (white) or 1% (gray) (w/w). FC= Field capacity. PWP=Permanent wilting point. Dashed lines represent fits to the van Genuchten model. (B) $CH_3Br$ standard curves in the same artificial soils. Artificial soils were hydrated to the same water content as used on each experimental design using MIDV1 media. Serial dilutions (10x) of $CH_3Br$ chemical standard were added to artificial soils containing OM.After capping, vials were incubated for 6 hours at 30°C prior to headspace gas measurements. Dashed line indicates a linear fit. Error bar indicates one standard deviation calculated using an n = 3.

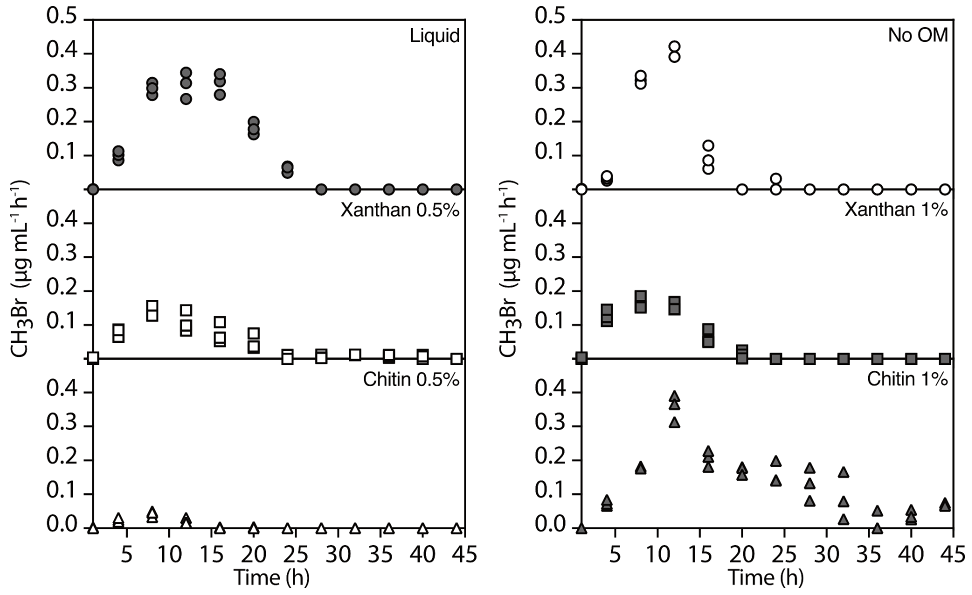


**Supplemental Figure 8. $CH_3Br$ production rates in soils containing OM.** $CH_3Br$ production per hour by Ec-MHT in artificial soils with different sources and amounts of OM, including xanthan (square) or chitin (triangle) at 0.5% (white) or 1% (gray) (w/w). An artificial soil without addition of OM, silt loam Q2a soil (white circle), and a liquid control (gray circle) are included. Each symbol indicates a single experiment. Experiments were performed in triplicates.

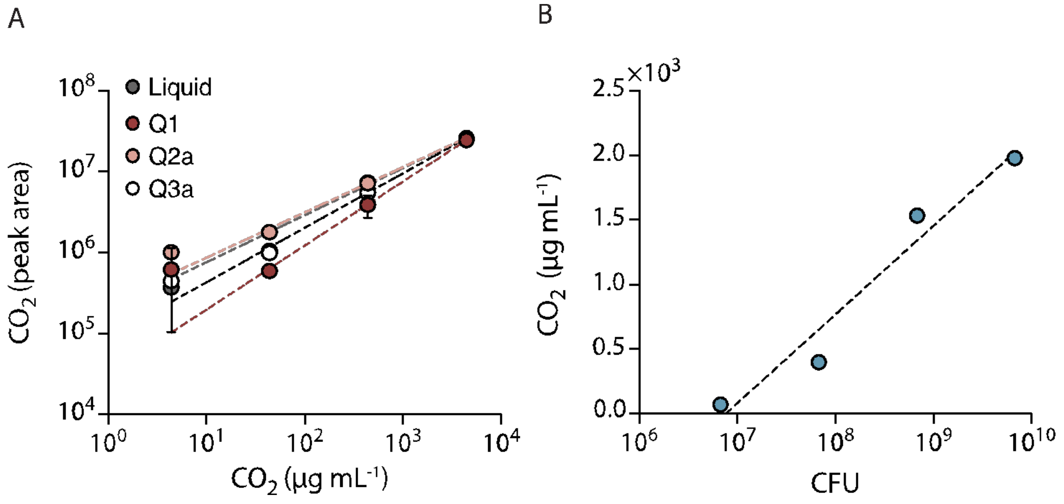


**Supplemental Figure 9. $CO_2$ standard curve and gas production per CFU.** (A) Serial dilutions (10x) of dissolved NaHCO3 were added to vials containing artificial soils with different particle size distribution (800 mg) or no matrix) in 100µL of MIDV1 medium. H3PO4 (100µL of 85%) was added to a final θ = 0.25 g(H2O)/g(soil), and vials were rapidly capped and incubated for 6 hours at 30°C. Headspace gas was quantified using GC-MS. Dashed lines represent log-log fits to the data with all soils presenting R2= 0.99. (B) Serial dilutions (10x) *Ec-AHL-MHT* in MIDV1 medium (200µL) were added to 2mL glass vials. Production of $CO_2$ after 3 hours at 30ºC was measured using GC-MS. Vials were then uncapped, cells were plated in LB-agar medium, and CFU were quantified after 24-hour incubation at 30°C. A fit of gas production per CFU fitted to a log-log linear regression model yielded a R2 = 0.90. Experiments were performed in triplicate, and error bars represent one standard deviation.

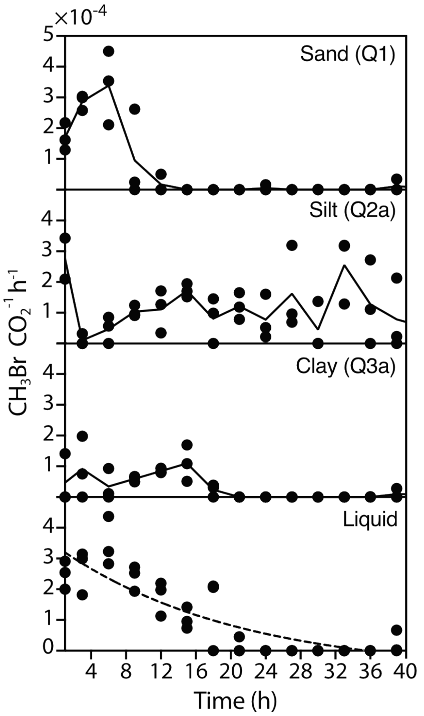


**Supplemental Figure 10. Particle size affects AHL-dependent $CH_3Br$ production**. $CH_3Br$ production rate after AHL addition (1 µM) to sand Q1, silt loam Q2a, clay Q3a, and liquid. Data was calculated using accumulation data shown in Figure 4. Each symbol indicates a different experiment. For each soil, the lines represent an interpolation of the mean values for each time. The dashed line for the liquid control indicates data fitted to a single exponential model (R2 = 0.73).

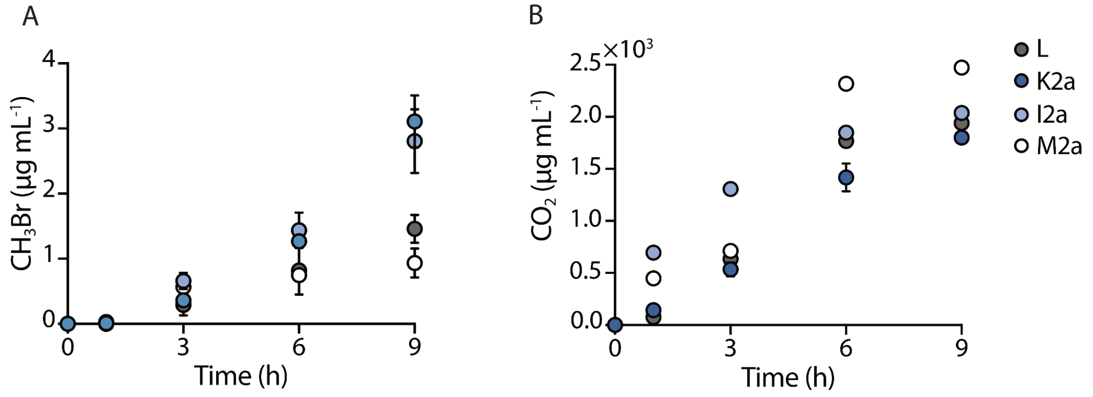


**Supplemental Figure 11. Constitutive gas production in soils with different mineralogies.** Ec-MHT (108 CFU) was incubated in soils with different mineralogy at ψm= -80kPa, and (A) $CH_3Br$ and (B) $CO_2$ accumulation was measured every 3 hours using a GC-MS. Experiment was performed in triplicates. Error bar indicates one standard deviation.

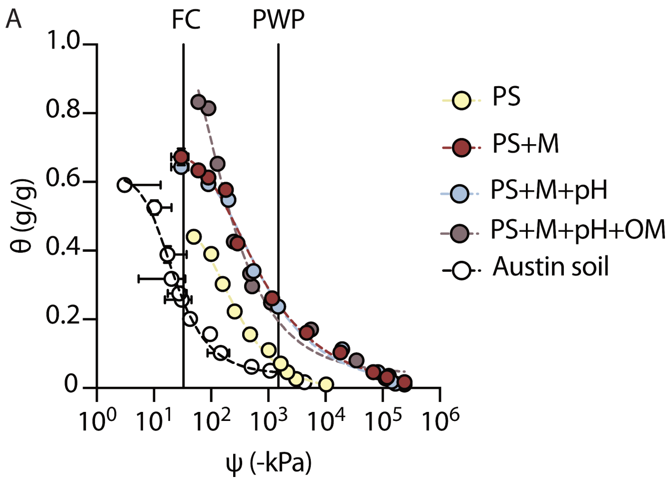


**Supplemental Figure 12. Water retention curve of the artificial soils built to mimic a Mollisol.** Water retention curve of the artificial soil series that recreate a natural Mollisol (Austin, TX) with increased complexity. PS = particle size, M = mineralogy, and pH = addition of $CaCO_3$, and OM = addition of xanthan gum. FC = field capacity and PWP = permanent wilting point. Error bars indicate one standard deviation calculated from three experiments.

**Supplemental Table 1. Fitted parameters of water retention curves.** Artificial soil water retention parameters calculated using the Van Genuchten model. Experimental data was analyzed using the SWRC fit program (Seki, 2007).

| **Van Genuchten model**<br>**$Se = [1/(1+(a*h)^n)]^m$**<br>**where $S_e = (\theta-\theta_r)/(\theta_s-\theta_r)$, $m=(1-1/n)$**<br>**h (meter) = ψ (Pa) / (ρw (1000 kg/m³) * g)** | | | | |
|---|---|---|---|---|
| **Soil Type** | **α** | **n** | **θ s** | **θ r** |
| Q1 | 0.091752 | 5.6762 | 0.22368 | 0 |
| Q2a | 0.086031 | 2.0222 | 0.41757 | 0.00019695 |
| Q3a | 0.041696 | 1.9634 | 0.44709 | 1.2371E-06 |
| K2a | 0.10613 | 1.6282 | 0.45243 | 3.2122E-06 |
| I2a | 0.1053 | 1.942 | 0.29701 | 0.001488 |
| M2a | 0.029284 | 1.8916 | 0.60725 | 0.009944 |
| Q3a-pH8 | 0.043909 | 2.5122 | 0.49741 | 0.01189 |
| Xanthan 0.5% | 0.33348 | 1.4425 | 0.60308 | 3.4269E-07 |
| Xanthan 1% | 0.019416 | 3.9543 | 0.77208 | 0.00002764 |
| Chitin 0.5% | 0.06104 | 1.9943 | 0.5143 | 0.015972 |
| Chitin 1% | 0.047132 | 2.7996 | 0.59148 | 0.036327 |
| Austin-PS | 0.092257 | 1.7244 | 0.48859 | 1.7085E-08 |
| Austin-PS+M | 0.07296 | 1.4289 | 0.69847 | 1.7454E-07 |
| Austin-PS+M+pH | 0.055626 | 1.4718 | 0.66416 | 0.0024949 |
| Austin-PS+M+pH+OM | 0.16333 | 1.5986 | 1.1087 | 0.03839 |
| Austin soil | 0.082429 | 1.9474 | 0.62327 | 0.039848 |

**Supplemental Table 2. Artificial soil pH values.** Soil pH was measured using a 1:1 water to soil ratio. Error indicates one standard deviation calculated using an n=3.

| Artificial soil | Average | error |
|---|---|---|
| Q1 | 7 | 0.01 |
| Q2a | 6.56 | 0.11 |
| Q3a | 7.15 | 0.18 |
| K2a | 7.9 | 0.09 |
| I2a | 8.65 | 0.06 |
| M2a | 9.2 | 0.11 |
| Q3a-pH8 | 8.51 | 0.01 |
| Xanthan 0.5% | 5.58 | 0.08 |
| Xanthan 1% | 5.39 | 0.07 |
| Chitin 0.5% | 5.65 | 0.02 |
| Chitin 1% | 5.66 | 0.02 |
| Austin-PS | 6.57 | 0.03 |
| Austin-PS+M | 7.63 | 0.10 |
| Austin-PS+M+pH | 8.42 | 0.04 |
| Austin-PS+M+pH+OM | 8.34 | 0.01 |

**Supplemental Table 3. Physicochemical properties of the Mollisol from Austin, TX.** Total nitrogen (N%), total soil carbon (TC%), organic carbon content (OC%), percentage of particle size (sand, silt, clay), surface area (SA), and pH of the natural soil from Austin, TX.

| Soil | N (%) | TC (%) | OC (%) | sand (%) | silt (%) | clay (%) | SA (m2/g) | pH |
|---|---|---|---|---|---|---|---|---|
| Average | 0.1 | 8.5 | 1.1 | 13.6 | 31.3 | 55.0 | 30.9 | 8.7 |
| Error | 0.0 | 0.1 | 0.1 | 0.5 | 1.0 | 0.5 | 0.8 | 0.08 |

**Supplementary information 2: Artificial Soils Protocol**

Adapted from *Artificial soils to analyze effects of matrix properties on microbial behaviors.*
Ilenne Del Valle, Xiaodong Gao, Teamrat Ghezzehei, Jonathan J. Silberg, and Caroline A. Masiello

**Materials**

- Sand-sized quartz grains (70 um) (NJ2 from U.S. Silica)
- Silt-sized quartz grains (~8.71 um) (Min-U-Sil 40 from U.S. Silica)
- Clay-sized quartz grains (~1.7 um) (Min-U-Sil 5 from U.S. Silica)
- Montmorillonite grains (CN: 18-600-868, Spectrum Chemical MFG Corp (Gardena, CA))
- Kaolinite grains (CN: 18-603-616, Spectrum Chemical MFG Corp (Gardena, CA))
- 250 mL plastic beaker
- Scoop
- Weigh paper
- 250 mL glass jar
- Additional glass jar (of desired size)
- DI water
- Spatulas
- Aluminum pan

**Equipment**

- Analytical balance
- Horizontal shaker (VWR)
- Oven
- 0.85 mm U.S. sieve
- 1.44 mm U.S. sieve

**Procedure**

The production of artificial soils requires the following steps 1) choose quartz grain size distribution, 2) add reactive minerals, if needed, 3) adjust soil pH, if needed, 4) add organic compound if needed, 5) create soil aggregation.

**Step 1: Texture**

1. Determine the mass and texture of artificial soil (AS) that will be needed. Calculate the appropriate mass proportions of each fraction of different sized quartz grain given in the USDA soil texture triangle (see below).

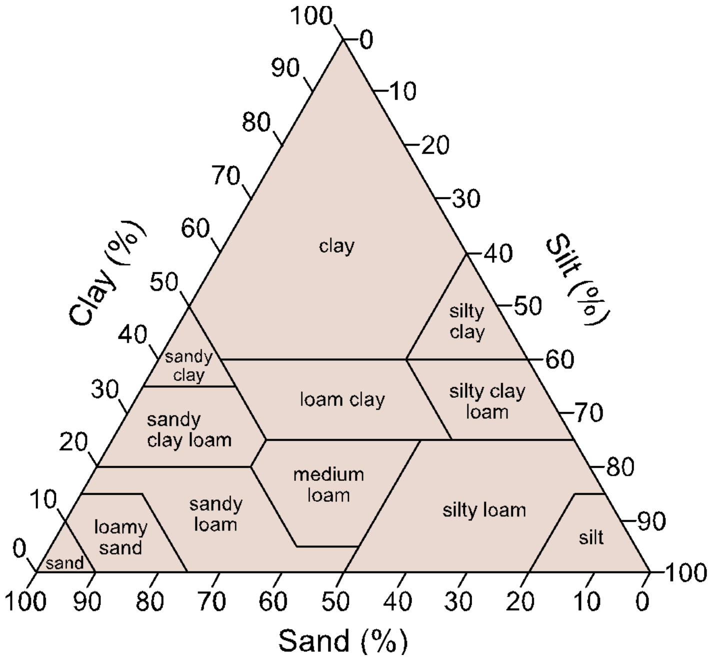


USDA soil texture triangle

2. Using the scoop, weigh paper, 250 mL plastic beaker and analytical balance, measure the quartz grains and record the exact mass of each component.
3. Pour all components into a 250 mL glass jar labeled with the AS type and cap the jar.
4. Manually mix the AS components in the jar for ~30 seconds. This can be done by shaking and repeatedly turning the jar.

5. Place the jar on a horizontal shaker at speed 7 rpm for 30 min. Remove the jar from the horizontal shaker.

**Step 2: Mineralogy**

1. If adding reactive minerals, replace the clay-sized quartz fraction for the mineral of interest in powder form.

> Note 1: Make sure that the reactive minerals are bought from a clean source (no organic or inorganic carbon contamination).
>
> Note 2: Soils with *sand* texture cannot be aggregated. Do not perform the following steps on artificial soils with these textures.

**Step 3: pH**

1. If adjusting soil pH (quartz-based particles make soil of roughly pH 7), $CaCO_3$ power (%w/w) can be added to increase soil pH as needed. More acid soils can be generated through the addition of $Al_2(SO_4)_3$ in powder (%w/w) to the dry soil mixture as needed.

**Step 4: Organic matter**

1. If adding organic matter, xanthan (surrogate for bacterial exopolysaccharides, contains C), chitin (surrogate for fungi-derived OM, contains C and N) or polygalacturonic acid (surrogate for plant mucilage) can be added in powder (%w/w) to the dry soil mixture as needed.

**Step 5: Aggregation**

1. Add Milli-Q water to the AS in its jar and stir constantly until all grains are hydrated and a slurry-paste consistency forms. *The amount of water required will vary with AS type.* The final consistency when wet will also vary with AS type, as indicated in Table 2:

**Table 2.** Consistency of each AS when wetted

| **Major component of AS** | Sand sized quartz | Silt sized quartz | Clay sized quartz | 1:1 Clay | 2:1 Clay |
|---|---|---|---|---|---|
| Final consistency | No wetting | Thick slurry | Slurry | Paste | Thick paste |

Note 3: For soils containing minerals with a 2:1 crystalline structure (expanding), soak mixed dry AS components in DI water for several minutes before stirring. This will help ensure even hydration of the grains and facilitate mixing.

2. Pour the wetted AS into a labeled aluminum pan and spread the paste to reach thickness of ~3-4 mm.
3. Place the pan in the oven to dry overnight at 60°C.
4. Rinse and dry labeled jar.
5. Remove from the oven and let it cool ~5 min. *This completes one wetting-drying cycle.*
5. Using a spatula, gently break up AS into pieces small enough to be placed in the original labeled glass jar.
6. Repeat steps 1-5 3 times to complete 3 additional wetting-drying cycles for a total of 4 wetting-drying cycles per AS.
7. After completing all the wetting-drying cycles, place the broken pieces of AS into a 2 mm sieve. Sieve the AS to 1-2 mm to create aggregates. Place in the original labeled jar and store for future use.

Note 4: AS that contain a large proportion of reactive minerals are harder and therefore more challenging to sieve without other tools. They can be gently broken in the 2 mm sieve with a spatula (1:1 clay) and/or pestle (2:1 clay).
